# Molecular Mechanisms of *Coxiella burnetii* Formalin Fixed Cellular Vaccine Reactogenicity

**DOI:** 10.1101/2024.08.20.608821

**Authors:** A. P. Fratzke, J. A. Szule, S. M. Butler, E. J. van Schaik, J. E. Samuel

## Abstract

Local and systemic reactogenic responses to Q-VAX® have prevented licensing of this vaccine outside of Australia. These reactogenic responses occur in previously sensitize individuals and have not been well defined at the cellular level, in part because many studies have been done in guinea pigs that have limited molecular tools. We previously characterized a mouse model of reactogenicity where local reactions sites showed an influx of CD8+ and IFNγ-expressing IL17a+ CD4+ T cells consistent with a Th1 delayed-type hypersensitivity. In this study we determined using depletion and adoptive transfer experiments that both anti-*Coxiella* antibodies and CD4+ T cells were essential for localized reactions at the site of vaccination. Furthermore, IFNγ depletion showed significant histological changes at the local reaction sites demonstrating the essential nature of this cytokine to reactogenicity. In addition to the cells and cytokines required for this response, we determined WCV material remained at the site of vaccination for at least 26 weeks post-injection. Transmission electron microscopy of these sites demonstrated intact rod-shaped bacteria at 2 weeks post-injection and partially degraded bacteria within macrophages at 26 weeks post-injection. Finally, since SCVs are an environmentally stable form, we determined that local reactions were more severe when the WCV material was prepared with higher levels of SCVs compared to typical WCV or with higher levels of LCV. These studies support the hypothesis that antigen persistence at the site of injection contributes to this reactogenicity and that anti-*Coxiella* antibodies, CD4+ T cells, and IFNγ each contribute to this process.

## Introduction

*Coxiella burnetii*, an obligate intracellular bacterium, is the causative agent of the zoonotic disease Q fever (1). This bacterium persists in the environment as a small cell variant (SCV) which is metabolically inactive and exceptionally resistance to heat and chemical disinfection (2). Transmission occurs after inhalation of contaminated aerosols or dust shed from infected animals (3). Once inhaled SCVs are phagocytosed by alveolar macrophages, they become metabolically active large cell variants (LCVs) after acidification of the phagolysosome, which is modified by effectors secreted by the T4SS to become the *Coxiella-*containing vacuole (CCV) (4). Exposure to this pathogen most often results in mild or asymptomatic infections, however Q fever can present as several severe chronic manifestations including myocarditis, placentitis, valvular endocarditis, and Q fever fatigue syndrome (5–8). Endemic worldwide, except New Zealand and Antarctica, this pathogen is maintained in environments through persistent zoonotic infections including in domestic ruminants and camelids (9). *C. burnetii* is an highly transmissible pathogen, with an infectious dose of 1-10 bacteria (3). This, in addition, to the highly resistant environmental SCV has resulted in *C. burnetii* being designated a Category B select agent by the Centers for Disease Control and Prevention (CDC) due to concerns for its potential as a weapon of bioterrorism. As such, a safe and effective vaccination program has been considered the best strategy to prevent infections in animal production industry and for military personnel stationed in endemic areas.

Q-VAX® is a phase I, formalin-fixed whole cell vaccine against Q fever and is licensed for use in humans in Australia. Vaccine protection is provided by both cellular and humoral components of the immune system in humans (10, 11). A retrospective study on the efficacy of Q-VAX reported that this vaccine provided a protective efficacy of 94.37% compared to unvaccinated (12). However, it has been associated with a high rate of local and systemic adverse responses which prevents its approval in other countries (13). These adverse reactions to Q-VAX are reported to be most severe and frequent in those with prior exposure to *C. burnetii* (14). To reduce the risk of these reactions, individuals undergo pre-vaccination screening that includes detection of anti-*C. burnetii* serologic titers and intradermal skin testing (15). Even though these reactions have been reported in humans and animal models for several decades, the mechanisms and causes for these vaccine-associated hypersensitivity reactions remain poorly defined (14). This is mainly because the hypersensitivity model of choice has been the guinea pig where immunological tools are limited. To overcome this limitation, we recently developed a mouse model of hypersensitivity, utilizing either post-vaccination or infection sensitization with *C. burnetii*, that demonstrated this response could be categorized as a Th1 delayed-type hypersensitivity response (16).

Histopathologic assessment of the vaccine site reactions to the *C. burnetii* formalin-fixed vaccine in humans and animal models showed pyogranulomatous and lymphocytic inflammation, which is characteristic of a granulomatous, delayed-type hypersensitivity (16–18). Delayed-type hypersensitivities are mediated by T cells, while the granulomatous component suggests that the inciting antigen is difficult to degrade or remove from the injection site (19). Some investigators developing novel vaccines against *C. burnetii* have begun to elucidate components of *C. burnetii* formalin-fixed whole cell vaccine (WCV) reactivity. For example, a recent study showed that vaccine reactions occur with both phase I and phase II derived WCVs in a sensitized guinea pig model, which we have also observed (22). Phase II *C. burnetii* is characterized by a large genetic deletion which eliminates O-antigen production resulting in rough LPS. This *C. burnetii* isolate, RSA 439, clone 4 is exempt from the select agent rule because it is attenuated (20, 21). This is significant because phase I LPS has been shown to be an important contributor to vaccine-mediated immunity with *C. burnetii* WCV, but did not significantly contribute to local reactogenicity, suggesting that difficult-to-remove antigens at the injection site were contributing to the response (22, 23). Furthermore, it was demonstrated that adjuvanted solubilized phase II material, unlike phase II WCV, was protective and no longer reactogenic, highlighting difficult-to-remove antigens contributing to the Th1-mediated hypersensitivity response (23, 24). The magnitude of the adaptive T cell response is also a contributing factor to the reactogenicity because an unadjuvanted subunit vaccine composed of six recombinant *C. burnetii* antigens was not reactogenic in sensitized guinea pigs, however, this vaccine showed moderate to severe reactions when mixed with a combination of squalene and toll-like receptor agonist adjuvants (16).

We previously investigated the immunopathogenesis of *C. burnetii* WCV reactions by characterizing the infiltrating immune cells in a sensitized mouse model (16). Immune cells extracted from vaccine site reactions showed an influx of CD4+ and CD8+ T cells. The CD4+ T cells were IFNγ+ and IL17α+, suggesting a Th1-mediated hypersensitivity reaction. However, the roles of these infiltrating T cells and cytokines in producing these local reactive lesions is still uncertain. Here we evaluated the roles of CD4+ and CD8+ T cells as well as the roles of IFNγ and IL17α in mediating *C. burnetii* WCV reactions at the injection site. Additionally, we show that injection of WCV leads to chronic persistence of antigen within the injection site which is likely contributing to the granulomatous component of local vaccine reactions. Furthermore, the severity of the response at the reactions site is more severe when the vaccine material is prepared by extending culture time to increase *C. burnetii* SCVs. This form of *C. burnetii* is a dense compact bacterial structure that is environmentally stable and produces more persistent antigen, however, the alternative is that antigens/structures present on SCVs that are not present on LCVs are the major contributor to Th1-mediated hypersensitivity at the injection site.

## Results

### CD4+ T Cells are essential for *C. burnetii* WCV hypersensitivity reactions

We previously determined that local vaccine reactions to *C. burnetii* WCV in sensitized mice is characterized by an increase in both CD4+ and CD8+ T cells (16). However, the specific roles of these T cell populations in mediating these hypersensitivity reactions were unknown. To evaluate the roles of CD4+ and CD8+ T cells in facilitating reactions to *C. burnetii* WCV in sensitized mice, we depleted either CD4+, CD8+, or both and elicitation sites were monitored for 14 days prior to collection for histopathology (Fig. 1A).

**FIG 1.**
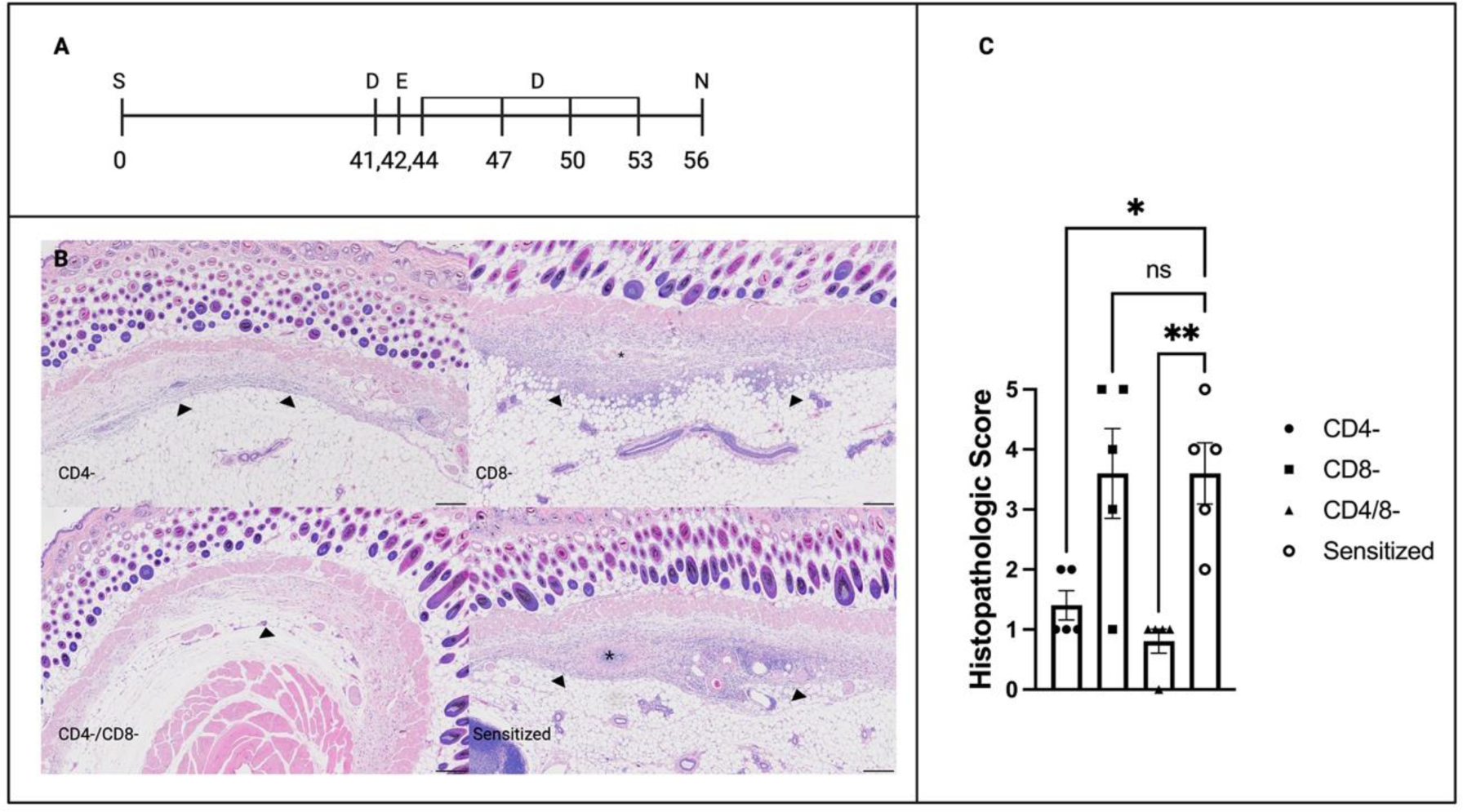
Histopathologic evaluation of vaccine site reactions in CD4 and CD8 T cell depleted mice. (A) Experimental time line. Each mouse was sensitized at time 0 and then antibody depletion was performed 1 day before elicitation and then 2, 5, 8, and 11 days post-elicitation. Necropsy was performed 2 weeks afer elicitation. A representative image of unsensitized control vaccine site is shown. (B) Representative images of vaccine sites reactions from each experimental group. Sensitized control mice and CD8-mice show marked lymphohistiocytic inflammation (arrows) with regions of suppurative necrosis (*). CD4- and CD4/8- mice show only mild lymphohistiocytic inflammation. HE-stained slides, 4x. (C) Semi-quantitative scoring of vaccine site reactions. CD4- and CD4/8- groups show a significant reduction in the severity of vaccine site reactions to C. burnetii WCV. N=5 mice per group. Significant difference are indicated with asterisks *, *P* < 0.05; **, *P* < 0.01; *** by one-way ANOVA with Dunnett’s correction for multiple comparisons.

HE-stained slides of vaccination sites showed a marked reduction in the severity of reactions in CD4- and CD4/CD8- groups compared to sensitized control mice based on semi-quantitative scoring of de-identified slides (Fig. 1B and C). Vaccine site reactions in CD8-mice did not reveal any significant changes in lesion severity or histomorphology compared to sensitized mice. Histology of vaccine sites from sensitized and CD8-mice showed granulomatous and lymphocytic inflammation with areas of suppurative necrosis and degeneration as well as dense aggregates of lymphocytes consistent with ectopic lymphoid follicles (Fig. 1B). In contrast, vaccine sites from CD4- and CD4/8- mice showed mild, dispersed histiocytic inflammation with small aggregates of lymphocytes (Fig. 1B). There was no evidence of suppurative necrosis or collagen degeneration in any tissue sections from the CD4- and CD4/8- groups. This indicated the CD4 T cells, but not CD8 T cells, are necessary to produce vaccine site hypersensitivity reactions.

### Adoptive transfer of serum enhances CD4+ T-mediated reactogenicity

We next evaluated whether local reactions to *C. burnetii* WCV reactions are mediated by memory CD4 T cell responses using adoptive transfer. Recipient mice were elicited with WCV as described in the previous experiment and vaccine site reactions were compared to sensitized and unsensitized controls (Fig. 2A).

**FIG 2.**
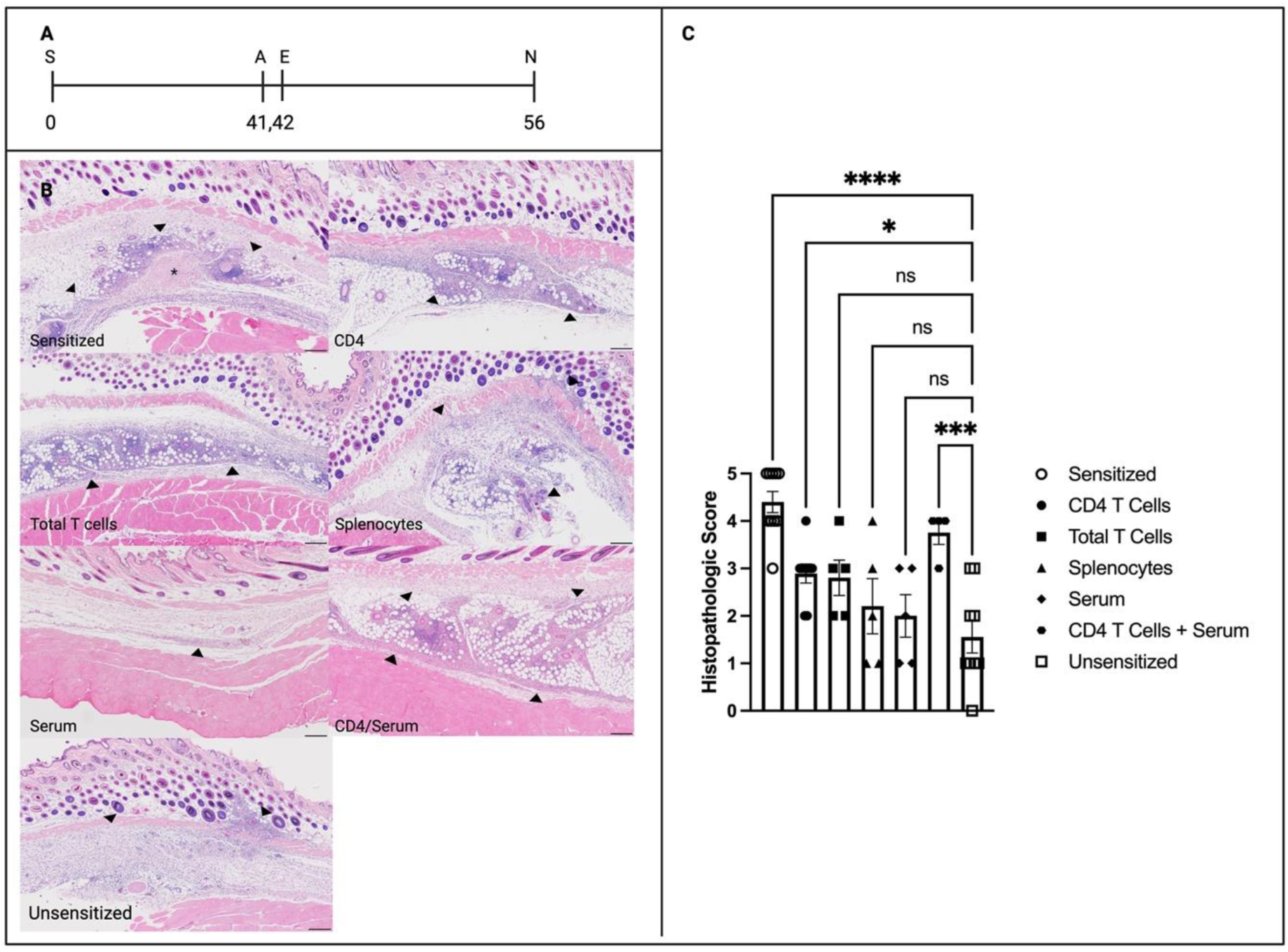
Histopathologic evaluation of vaccine site reactions in adoptively and passively transferred mice. (**A**) Experimental time line. Each mouse was sensitized at time 0 and then adoptive transfer was performed 1 day before elicitation. Necropsy was performed 14 days after elicitation. (**B**) Representative images of vaccine site reactions from experimental groups. HE, 4x, bar = 200 µm. (**C**) Semi-quantitative scoring of vaccine site reactions. CD4 T cell and CD4 T cell plus serum groups developed significantly more severe vaccine site reactions than unsensitized mice. Only CD4 T cell plus serum recipient mice show vaccine site reactions which do not significantly differ in severity from sensitized mice. N=5-10 mice per group. Significant difference are indicated with asterisks *, *P* < 0.05; **, *P* < 0.01; ***, *P* < 0.001; and ****, *P* < 0.0001 by one-way ANOVA with Dunnett’s correction for multiple comparisons.

Mice receiving either CD4 T cells or a combination of CD4 T cells and serum showed significantly more severe vaccine site reactions compared to unsensitized mice based on semi-quantitative scoring (Fig. 2B). However, the CD4 T cell only group also had less severe inflammation compared to the sensitized group. Morphologically, both the CD4 T cell and CD4 T cell plus serum groups showed lymphohistiocytic inflammation and formation of ectopic lymphoid follicles, but CD4 T cell plus serum group showed consistently more severe inflammatory cell infiltrates compared to the CD4 T cell group (Fig. 2C). None of the reaction sites from these two groups contained areas of suppurative necrosis as observed in the sensitized control group. Total T cell and splenocyte groups also showed mild to moderate lymphohistiocytic inflammation, but these lesions were less severe than sensitized mice. The serum recipient group show no apparent increase in the severity of reactive lesions compared to unsensitized mice. These results indicated that although anti-*C. burnetii* antibodies alone do not produce hypersensitivity reactions to *C. burnetii* WCV, they appear to enhance reactive lesions produced by CD4 T cells.

### Depletion of IFNγ alters local vaccine site reactions

Our previous work showed that infiltrating IFNγ+ and IL17α+ CD4 T cells extracted from elicitation sites are significantly increased in sensitized mice compared to unsensitized controls (13). To investigate how CD4 T cells may mediate local vaccine site reactions, we assessed the role of IFNγ and IL17α in producing hypersensitivity responses using depletion studies. Histology of the vaccine sites from sensitized and IL17α-mice were morphologically similar, showing severe lymphocytic and granulomatous inflammation with central areas of suppurative necrosis (Fig. 3A). Histology of vaccine sites in IFNγ- and IFNγ/IL17α-mice showed lymphohistiocytic inflammation as seen in sensitized controls, but lacked areas of suppurative necrosis. However, the overall severity of the vaccine sites reactions in depleted mice compared to sensitized mice did not significantly differ (Fig. 3B). To better assess these changes in morphology, semi-quantitative scoring of the inflammation was separated into three categories: suppurative necrosis, immune cell infiltration, and ectopic lymphoid follicles (Table 2, Fig. 3B). None of the vaccine site reactions in IFNγ- and IFNγ/IL17α-showed evidence of suppurative necrosis, however, these groups showed a significant increase in the severity of immune cell infiltrate compared to sensitized mice. Additionally, IFNγ- and IFNγ-/IL17α-mice showed a significant increase in number of ectopic lymphoid follicles compared to unsensitized mice. Morphologically, in sensitized mice the infiltrating immune cells were more dispersed and have a moderate amount of eosinophilic cytoplasm with distinct cytoplasmic borders, consistent with activated macrophages. In contrast, the immune cell infiltrate from IFNγ- and IFNγ/IL17α-mice had abundant cytoplasm with indistinct cell borders (Fig. 3C). Flow cytometry of cells extracted from vaccine site reactions was performed to quantify infiltrating cell numbers. Although not statistically significant, CD45+CD11b+Ly6G-macrophages were decreased in IFNγ- and IFNγ/IL17α-mice compared to sensitized mice (Fig. 4). Similarly, numbers of CD45+CD11b+Ly6G+ neutrophils were mildly decreased in all three depletion groups compared to sensitized mice.

**FIG 3.**
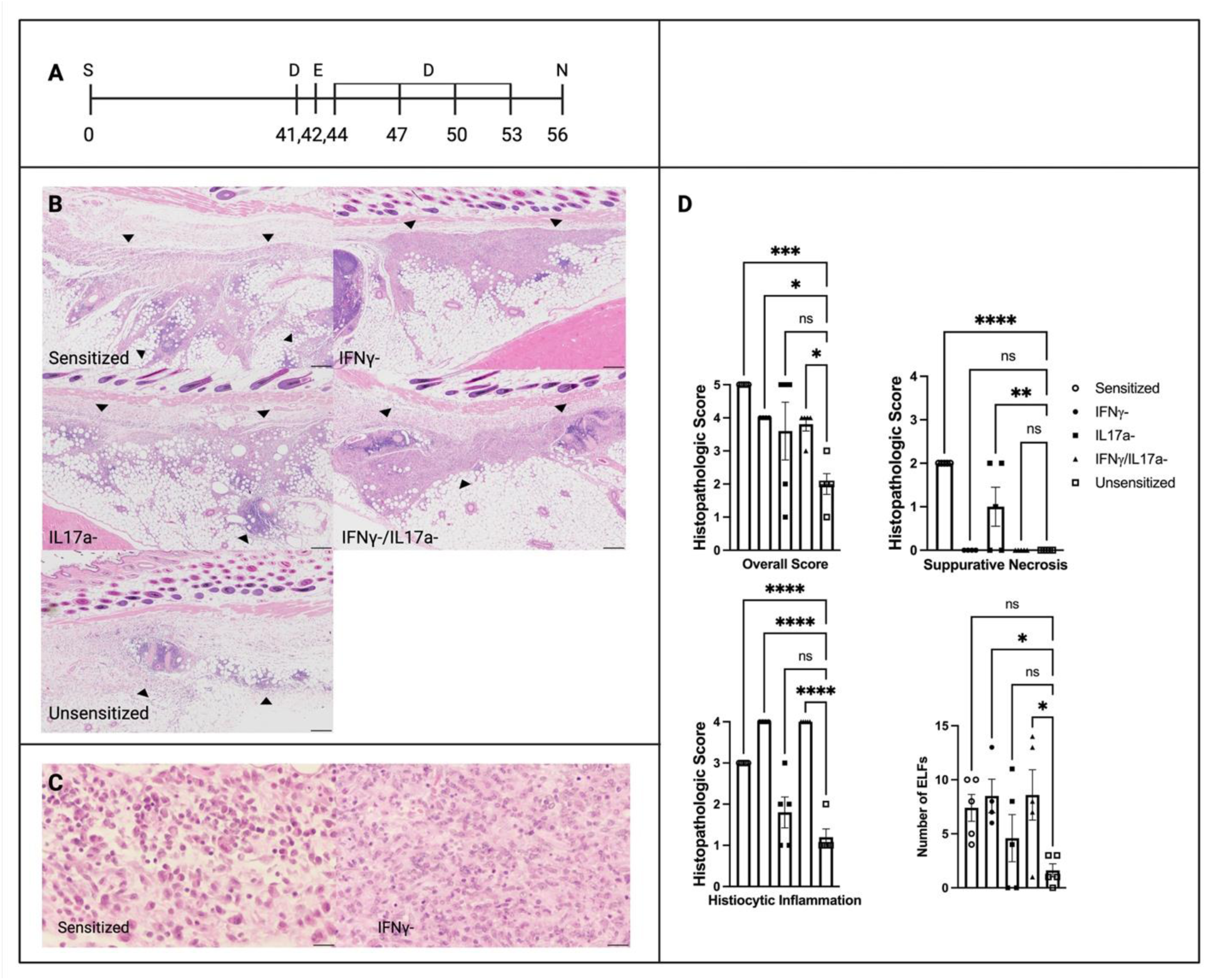
Histopathologic evaluation of vaccine site reactions in cytokine depleted mice. (**A**) Experimental time line. Each mouse was sensitized at time 0 and then antibody depletion was performed 1 day before elicitation and then 2, 5, 8, and 11 days post-elicitation. (**B**) Representative images of vaccine site reactions from experimental groups. HE, 4x, bar = 200 µm. (**C**) Vaccine site reactions in IFNγ-depleted mice showed dense infiltration of innate immune cells compared to the dispersed immune cells with distinct cell borders seen in sensitized control mice HE, 40x, bar = 25 µm. N=5-10 mice per group. (**D**) Semi-quantitative scoring of vaccine site reactions. Cytokine-depleted mice did not have significantly less severe reactions overall compared to sensitized controls. IFNγ-depleted mice showed a lack of suppurative necrosis, but an increase in immune cell infiltration compared to sensitized mice. Significant differences are indicated with asterisks *, *P* < 0.05; **, *P* < 0.01; ***, *P* < 0.001; and ****, *P* < 0.0001 by one-way ANOVA with Dunnett’s correction for multiple comparisons.

**FIG 4.**
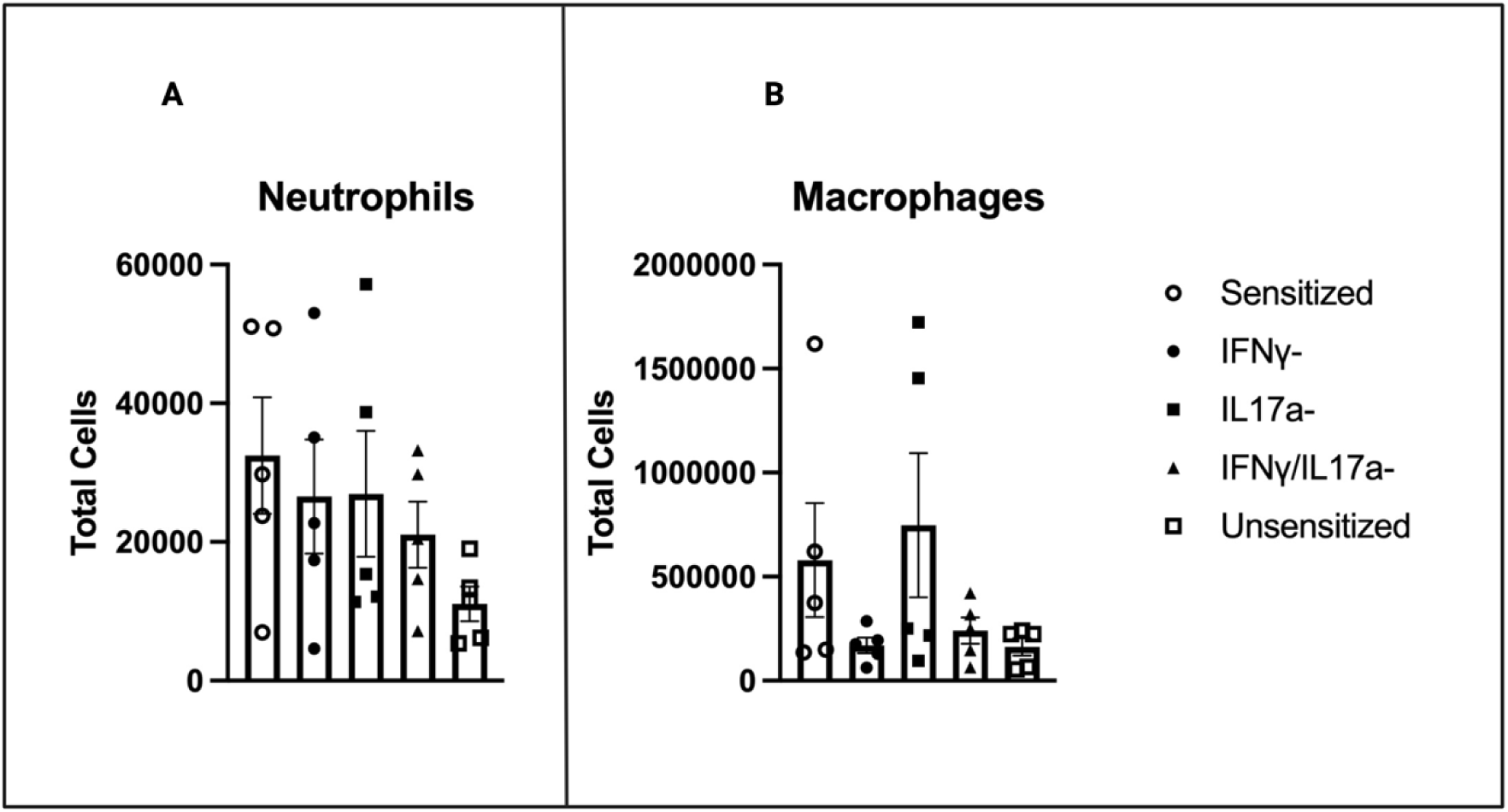
Flow cytometry of cells extracted from skin sections at the vaccine site. Summary graphs of total (**A**) CD11b+CD11c-Ly6G+ neutrophils and (**B**) CD11b+CD11c-Ly6G-macrophages. All depletion groups and unsensitized mice show a mild decrease in neutrophils compared to sensitized mice. IFNγ- and IFNγ-/IL17a- mice show a decrease in macrophages compared to sensitized mice. N=5 mice per group.

### Antigens from the C. burnetii WCV persist within injection sites for extended periods

The granulomatous component to the vaccine site reactions with *C. burnetii* WCV suggests an antigen which is difficult to degrade or remove from the injection site. To investigate how persistence of antigen may be contributing to vaccine site inflammation we evaluated areas of injection site reactions were evaluated at 8 weeks and 26 weeks post-injection. IHC with a polyclonal anti-*C. burnetii* Nine Mile I (NMI) antibody showed positive staining, usually within central regions of suppurative necrosis and within the cytoplasm of adjacent macrophages, at all evaluated time points (Fig. 5A).

**FIG 5.**
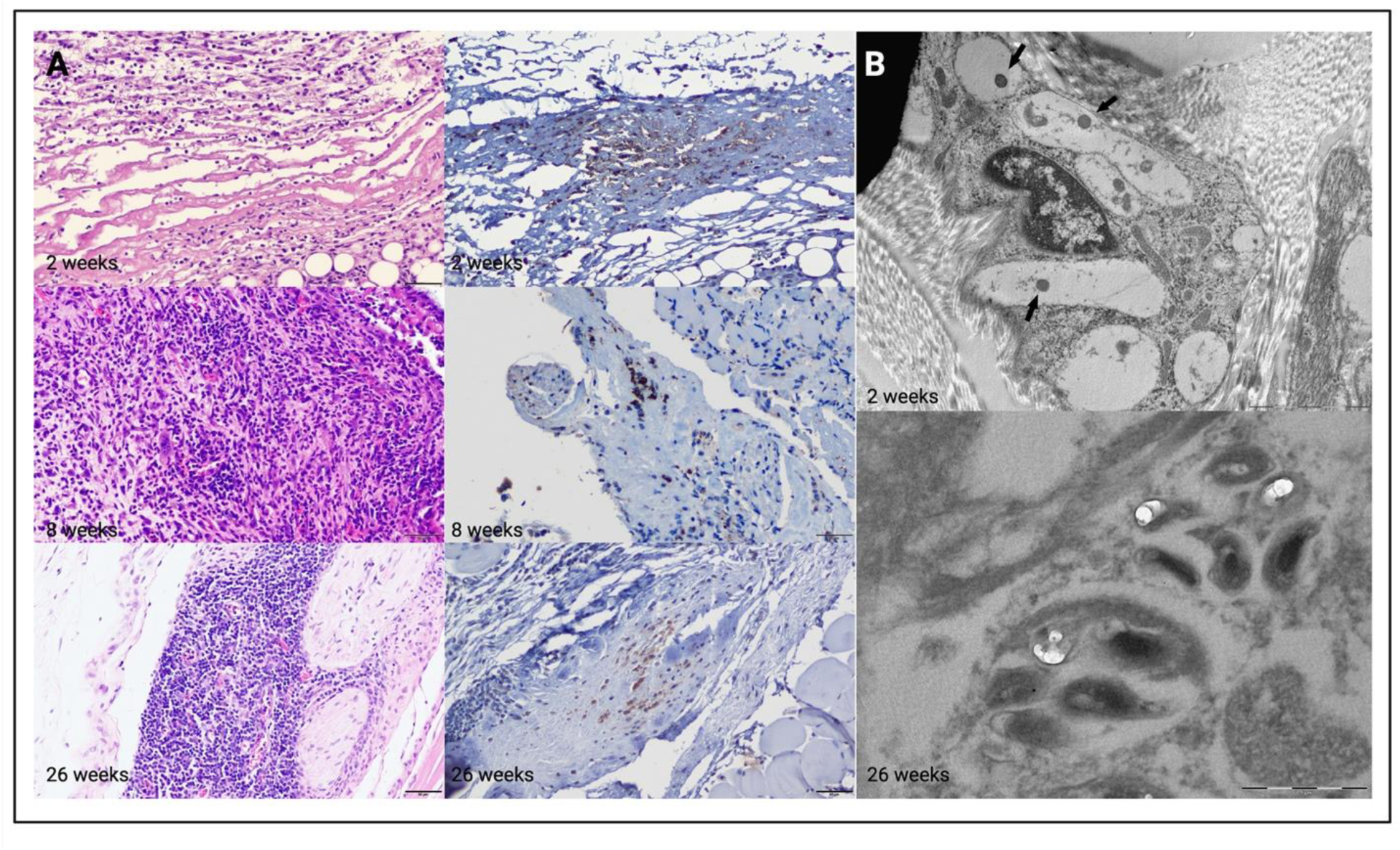
Histopathology, immunohistochemistry and transmission electron microscopy of vaccine sites in mice from different time points. (**A**) Representative images of vaccine site reactions with WCV at 2 weeks (10 µg dose), 8 weeks and 26 weeks (50 µg dose). Anti-*C. burnetii* IHC shows positive immunostaining at all time points. HE and DAB with hematoxylin counterstain, 20x, bar = 50 µm. Panels go from 2 weeks at the top to 26 weeks at the bottome with H&E on the left and αC.b. on the right (**C**) TEM of a vaccine site at 2 weeks and 26 weeks post-injection with WCV. Images from 2 weeks show coccobacilli measuring up to 500 nm in length. At 26 weeks, homogeneous variably shaped inclusions (partially degraded C. burnetii) are evident in vacuoles within macrophages arrows. Bars = 1 µm, 500 nm, and 100 nm.

To further investigate the etiology of this persistent antigen, we collected regions of tissues from paraffin-embedded tissue blocks which were stained positively with anti-*C. burnetii* antibodies to evaluate using transmission electron microscopy (TEM). TEM of the anti-*C. burnetii* positive region from an elicitation site at 2 weeks post-injection revealed numerous, apparently intact, bacteria (Fig. 5B). These bacteria were rod-shaped, measuring 0.2 to 0.3 µm long and 0.1 to 0.2 µm wide, with a dense, central nucleoid. At 26 weeks post-injection, homogeneous, variably shaped inclusions were evident in macrophages within the injection (Figure 5B). The size and shape of these inclusions are similar to the intact coccobacilli observed at the 2 week old site, indicating that these inclusions are partially degraded *C. burnetii*. This indicates that slow degradation and removal of the intact bacteria by host cells *in vivo* is the cause of local antigen persistence.

To further investigate the slow degradation and persistence of antigen, we created vaccine material for cultures that should be dominated by LCV or SCV to determine if the SCV form of *C. burnetii* would also cause more severe reactogenicity (Fig. 6). Histopathologic evaluation of injection sites showed lymphohistiocytic inflammation with central regions of suppurative necrosis and ELF formation in mice elicited with WCV, SCV and LCV. However, suppurative necrosis was more consistently observed in injections in mice elicited with SCV than LCV or WCV. These areas of suppurative necrosis are where persistent Cb antigen was observed in our IHC and TEM experiments, supporting the concept of SCV causing local persistence of antigen and significantly contributing to one of the histopathologic features of WCV reactogenicity.

**FIG 6.**
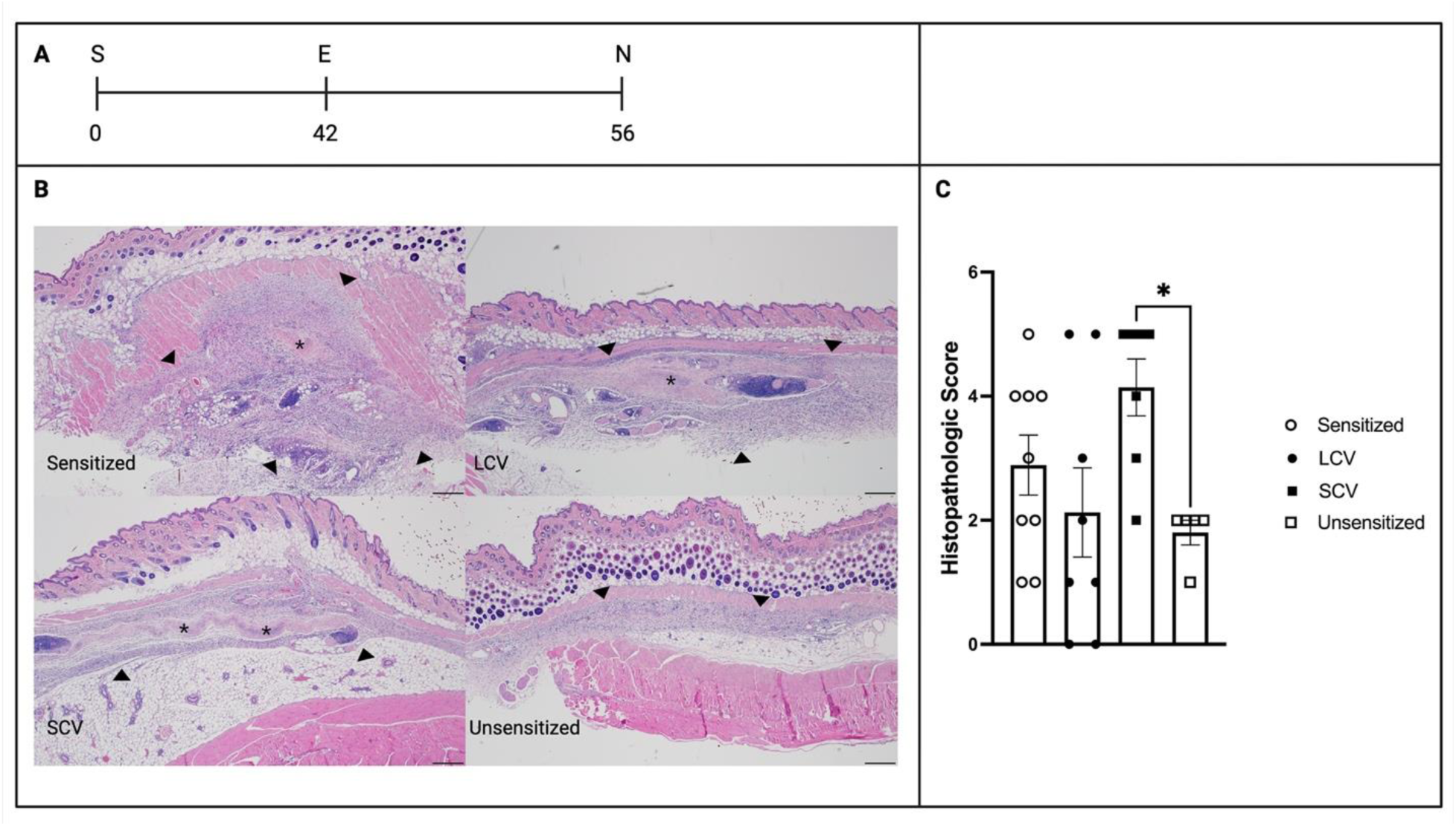
Histopathology evaluation of mice after stimulation with WCV material prepared from LCV and SCV dominant cultures. (**A**) Representative images of vaccine sites reactions from each experimental group. Histopathology of injection sites from sensitized control mice and SCV- and LCV-elicited mice have similar reactions consisting of lymphohistiocytic inflammation with central regions of suppurative necrosis. (**B**) Semi-quantitative scoring of vaccine site reactions. N=5 mice per group. Significant differences are indicated with asterisks*, *P* < 0.05 by one-way ANOVA with Dunnett’s correction for multiple comparisons.

## Discussion

While the severe reactogenic responses produced by vaccines against *C. burnetii* have long stood as a major barrier to the widespread availability of a protective vaccine, the mechanisms and causes underlying these reactions are still poorly understood. It was previously shown that local *C. burnetii* WCV reactions are characterized by an influx of CD4+ and CD8+ T cells in a sensitized mouse model (16). To investigate the roles of these infiltrating T cells in mediating *C. burnetii* WCV reactogenicity, we performed antibody-mediated depletion and adoptive transfer experiments in a sensitized mouse model. Depletion of CD4, but not CD8 T cells in sensitized mice prior to elicitation with WCV markedly reduced the severity of local inflammation compared to control mice, indicating that memory CD4 T cells play an essential role in producing *C. burnetii* WCV reactogenicity (Fig. 1). However, adoptive transfer of immune CD4 T cells alone did not fully re-create the severity of local vaccine reactions in sensitized mice. Interestingly, the addition of immune serum with adoptive transfer of memory CD4 T cells enhanced the severity of local reactions (Fig. 2). This indicated that although local reactions are mediated by memory CD4 T cells, their activity is enhanced by a component of immune serum, presumably anti-*C. burnetii* antibodies (Fig. 2).

Prior studies in contact and delayed-type hypersensitivities have already indicated a role for antigen-specific antibodies in T cell-mediated reactions. Antigen-specific IgM produced by immune B1 cells binds to antigens and activates complement, leading to T cell recruitment during elicitation (16). Anti-*C. burnetii* antibodies and more specifically anti-phase I LPS antibodies may similarly be enhancing CD4 T cell recruitment during elicitation with WCV. In previous work, antibodies targeting the phase I LPS of *C. burnetii* have been shown to play an important role in the protective efficacy of the *C. burnetii* WCV (23, 25). Anti-LPS antibodies may similarly be enhancing immune responses to WCV during elicitation, leading to more severe local reactions. However, further experiments are needed to elucidate the roles of anti-*C. burnetii* antibodies in mediating *C. burnetii* WCV reactogenicity and whether both LPS and protein specific antibodies contribute to this response. Although both WCV from phase I and phase II *C. burnetii* cause reactogenicity, the response to phase II is more variable, which suggests that anti-LPS antibodies could still play a role only when CD4 T cells and protein antigens are present (22, 26).

Notable here is that both memory CD4 T cells and anti-*C. burnetii* antibodies have been shown to be important for WCV-meditated protection during infection (23, 25). WCV elicits a Th1-dominant immune response which leads to formation of memory T cells and antibodies specific for phase I LPS which are essential for WCV-mediated protection (23). Thus, the mechanisms of WCV-mediated protection appear to parallel the mechanisms of local reactogenicity. This, however, does not necessarily mean that severe reactogenicity is inevitable with an efficacious vaccine against *C. burnetii*. Studies on immune responses during *C. burnetii* infection indicate that although T cells are necessary for control of *C. burnetii* during infection, CD8 T cells, which do not appear to contribute to reactogenicity, play an important role in clearance of bacteria during infection (27, 28). Additionally, mice depleted of CD4 T cells not only show decreased bacterial load, they display less lung pathology during pulmonary challenge with *C. burnetii*, suggesting that reducing CD4 T cell responses will not preclude protection (27).

We next investigated the roles of IFNγ and IL17α in producing local *C. burnetii* WCV reactions. Previously, these cytokines were shown to be increased during elicitation of hypersensitivity reactions to WCV (29). Depletion of IFNγ and IL17α did not markedly reduce the overall severity of local hypersensitivity reactions as seen when mice were depleted of CD4 T cells and mice depleted of IL17α showed no appreciable changes in the morphology of hypersensitivity reactions compared to sensitized mice (Fig. 1 and 3). This was surprising considering that abscesses are commonly seen in *C. burnetii* WCV reactions and IL17 is an important activator of neutrophils which has been implicated in abscess formation (30). However, in this experiment we used antibodies specific for IL17α to deplete mice. Although IL17α is considered the main inducer of neutrophil responses in contact and delayed-type hypersensitivity, IL17f has also been shown to be an important mediator of neutrophil responses during bacterial infection (31). Since *C. burnetii* WCV reactogenicity is caused by responses to bacterial antigens, it is possible that IL17f or another IL17 subtype is involved in mediating neutrophil responses in *C. burnetii* WCV reactogenicity.

In contrast to IL17α-depleted mice, IFNγ-depleted mice showed significant changes in the morphology of local reactions. Histologically, the local reactions in IFNγ-depleted mice lacked regions of suppurative necrosis and inflammation was composed of dense immune cell infiltrates with indistinct cytoplasmic borders, while sensitized mice show dispersed immune cells with moderate cytoplasm and distinct cell borders. IFNγ is an important inducer of macrophage priming and activation. Classically activated macrophages, or M1 macrophages, are stimulated by a combination of IFNγ and TLR signaling leading to altered cellular metabolism which enhances pro-inflammatory cytokine production and phagocytic activity (32). The lack of IFNγ in depleted mice may be inhibiting normal macrophage activation and phagocytosis of antigens, leading to the altered morphology of the inflammatory infiltrate. For the absence of suppurative necrosis, CD4 T cells, especially Th1 cells, are known to be important in the formation of abscesses, and IFNγ production by CD4 and CD8 T cells enhances neutrophil activation (30, 32, 33). Although IFNγ appears to play a significant role in mediating *C. burnetii* WCV reactions, the presence of severe local reactions despite depletion of IFNγ indicates that CD4 T cells mediate these reactions by other mechanisms as well.

Lastly, we showed that the *C. burnetii* WCV causes prolonged persistence of antigen at the injection site (Fig. 5). Granulomatous delayed-type hypersensitivity reactions are associated with an antigen which is difficult to degrade or remove (19). IHC for *C. burnetii* on vaccine sites showed positive immunostaining at 2 weeks, 8 weeks, and 26 weeks post-injection (Fig. 5). TEM of the 2 week and 26 week post-injection sites revealed intact coccobacilli and partially degraded remnants of *C. burnetii* within macrophages, respectively (Fig. 5). The inability of host immune cells to degrade and remove intact *C. burnetii* from the injection is very likely a significant contributor to chronic reactogenicity observed with WCV.

The inability to remove antigen suggested that SCVs in WCV may cause more severe reactogenic responses than LCVs since it is the stable environmental form which resists degradation. Indeed, vaccine material prepared from LCV- and SCV-enriched cultures confirmed that SCVs caused more severe reactions than LCVs and WCV (Fig. 6). However, another contributing factor to the more severe reactogenicity of SCV could be different structural commponents. Furthermore, as it was recently demonstrated that the type IV secretion (T4SS) apparatus was absent from SCVs (4), our data suggest that the T4SS is not the contributor of reactogenicity in SCVs, although others have shown the T4SS can contribute to this response in WCV material (22). Additionally, formalin-fixation, which causes cross-linking of proteins, is used to produce *C. burnetii* WCV and Q-VAX and may produce a depot effect, reducing the rate of clearance of antigen from the injection site, which can contribute to both protective efficacy and reactogenicity of these vaccines (34). Therefore, preparation of a less persistent vaccine material will likely reduce local reactogenicity as we previously demonstrated with phase II adjuvanted soluble material (24). Our current data suggest that the potential to make vaccine material from LCVs using an inactivation process that does not inhibit degradation of antigens, like gamma-irradiation or solubilization could lead to a vaccine that is protective and non-reactogenic (Fig. 5) (35, 36). Others have reported that vaccines derived from individual proteins or extracts of *C. burnetii* are able to reduce the severity of local reactogenicity (24, 37, 38). A similar reduction in vaccine reactogenicity has also been described when comparing whole cell and acellular pertussis vaccines (39). The mechanism behind reduced reactogenicity when comparing cellular versus acellular vaccines may be due to a reduction in local antigen persistance.

Differentiation of protective and pathologic mechanisms of vaccine-induced immune memory is essential for development of safer vaccines. As discussed above, CD4 T cells, immune serum, IFNγ, and local persistence of antigen all contribute to *C. burnetii* WCV reactogenicity. Use of subunit or whole cell vaccines without factors that may inhibit antigen degradation, such as formalin and SCVs, will likely dramatically reduce the severity of local persistence of antigen. Indeed, the injection of unadjuvanted *C. burnetii* antigens does not appear to induce significant reactogenicity (37). However, this creates a significant reduction in protective efficacy, necessitating the use of adjuvants which can also cause marked reactogenicity when paired with *C. burnetii* antigens (37). Knowing that reactogenicity with WCV is mediated by CD4 T cells, adjuvant combinations in vaccines should be tailored towards producing CD8 rather than CD4 T cell memory responses. Using a combination of several TLR agonists, including TLR4 agonists, and Addavax, a squalene-based oil-in-water emulsion demonstrated that only one of five tested formulations showed a significant reduction in reactogenicity compared to sensitized animals (37). Addavax stimulates innate immune cells to create an immunocompetent environment which enhances CD4 T cell and B cell responses (40). Similarly, TLR4 agonists are strong inducers of CD4 T cell memory (41). Together the effects of these adjuvant combinations were likely producing strong CD4 T cell memory and antibody responses which could be the cause of the severe local reactions reported. In contrast, addition of a TLR9 agonist, CpG ODN 1826, to solubilized protein extract from *C. burnetii* not only made normally non-protective material protective, it also reduced the reactogenicity (24). TLR9 is a strong inducer of CD8 T cell memory, which may partially explain the reduction in reactogenicity with the preservation of protection reported in those experiments (42). Overall, reduction in reactogenicity with novel *C. burnetii* vaccines will likely require two main strategies. First, reducing local persistence of antigen to minimize granulomatous inflammation, and second, a focus on induction of strong memory CD8 T cell responses rather than CD4 T cell responses to reduce pathologic adaptive responses.

Here we investigated the roles of T cells, IFNγ, and IL17α in mediating *C. burnetii* WCV reactogenicity as well as the long-term persistence of antigen from *C. burnetii* WCV. Local vaccine site reactions in sensitized mice are mediated by CD4 T cells and enhanced by a component of immune serum, presumably anti-*C. burnetii* antibodies and potentially anti-LPS antibodies in addition to anti-*Coxiella* protein antibodies. Additionally, IFNγ plays a significant role in the morphology of local reactive lesions, but is not required for induction of the reaction. Lastly, SC injection of *C. burnetii* WCV causes localized persistence of antigen which is likely contributing to the chronicity of local inflammation and SCV-enriched vaccine material increased the severity of reactogenic response. Our work provides insights into the mechanisms of *C. burnetii* WCV reactogenicity which may help guide the development of novel protective vaccines with enhanced safety profiles.

## Materials and methods

### Vaccine Materials

WCV was produced by growing cultures of *C. burnetii* Nine Mile Phase I (NMI) RSA493 in ACCM-2 media as previously described, then inactivated in 2% formalin for 48 hours (37, 43). LCV- and SCV-enriched preparations were cultured for 5 and 28 days, respectively prior to formalin-inactivation (44). Culture of live *C. burnetii* NMI RSA493 was performed in biosafety level 3 (BSL3) facilities at the Texas A&M School of Medicine.

### Experimental animals

Female C57Bl/6JHsd mice at 6-8 weeks old were purchased from Envigo (Huntingdon, UK). Mice were housed in microisolator cages under pathogen-free conditions with ad-lib food and water. Animals were housed in either biosafety level 1 or level 2 rooms and all experiments were performed with an approved animal use protocol as reviewed by the Institutional Animal Care and Use Committee at Texas A&M University.

### Sensitization and Elicitation of Responses

For sensitization, mice were injected subcutaneously (SC) with 50 µg of WCV in 50 µL sterile PBS in the middle of the back, then rested for 6 weeks prior to elicitation. Mice were elicited with WCV as previously described (16). Briefly, mice were anesthetized by IP injection of 100 mg/kg ketamine and 10 mg/kg xylazine. Hair was removed from elicitation sites using electric clippers followed by application of a depilatory cream. Mice were vaccinated SC with 10 µg WCV and elicitation sites were monitored for 14 days prior to collection for histopathology or flow cytometry.

### Antibody-Mediated Depletion

For antibody-mediated depletion experiments, mice were injected IP with 200 µg anti-mouse antibodies 1 day prior to elicitation, followed by 150 µg at days 2, 5, 8, and 11 post-elicitation. Antibodies include anti-CD3ε (GK1.5), anti-CD8 (2.43), anti-IFNγ (XMG1.2) (45), anti-IL17α(17F3) (46), and isotype control (LTF-2) from BioXCell (Lebanon, NH, USA). Depletion of cells was evaluated by collecting cells from spleens and vaccine sites of depleted mice and staining with anti-CD3ε, anti-CD4, and anti-CD8a (Biolegend) (Supplemental material).

### Adoptive and Passive Transfer

Donor mice were sensitized as previously described and rested for 6 weeks prior to collection. Spleens and axillary lymph nodes were collected from donor mice and formed into a single cell suspension by pressing through a 70 µm cell strainer, twice in flow cytometry staining buffer (FACs buffer). Splenocytes were treated with ACK lysis buffer for 1 min to remove red blood cells. CD4 4+ T cells and total T cells were purified from cell suspensions using Mouse CD4+ T cell and total T cell Isolation Kits from Miltenyi Biotec (Auburn, CA, USA) (Supplemental material). Mice in each recipient group received a retro-orbital injection of 5×10^6^ cells in 50 µL sterile PBS or PBS alone one day prior to elicitation. Aliquots of purified cells were evaluated on flow cytometry to assess purity of transferred samples. Blood was collected from donor mice at euthanasia and used to provide serum for passive transfer. An aliquot of this serum measured anti-*C. burnetii* antibody titers of >3200 by ELISA performed as previously described (29).

### Histopathology and Immunohistochemistry

At 14 days post-elicitation, vaccine sites (skin, subcutis, and underlying skeletal muscle) were collected entirely and placed in 10% neutral buffered formalin for a minimum of 24 hours. Tissues were serially trimmed (3-4 sections per tissue) and placed in cassettes before being sent for processing, embedding, and sectioning at 5 µm (AML Laboratories, Jacksonville, FL, USA). Slides were stained with hematoxylin and eosin (HE) for assessment by a board-certified pathologist. Slides were de-identified and scored for overall severity based on lesion size, immune cell infiltrate, and regions of suppurative necrosis (Table 1) or separately evaluated for major morphologic features: histiocytic inflammation, suppurative necrosis, and formation of ectopic lymphoid follicles (Table 2). For immunohistochemistry, 5 µm sections were cut from tissue blocks and placed on adhesive slides then processed as previously described (29). Primary antibodies used for immunohistochemistry include rabbit anti-*C. burnetii* polyclonal serum at 1:500.

**TABLE 1.** Semi-quantitative scoring system for evaluation of histopathology of vaccine site reactions on HE-stained slides.

| Overall Histopathologic Score |  |
| --- | --- |
| 0 | No lesions |
| 1 | Minimal lymphohistiocytic inflammation |
| 2 | Mild, multifocal lymphohisitocytic inflammation |
| 3 | Moderate lymphohistiocytic inflammation |
| 4 | Severe lymphohistiocytic inflammation |
| 5 | Severe lymphohistiocytic inflammation with suppurative necrosis |

**TABLE 2.**
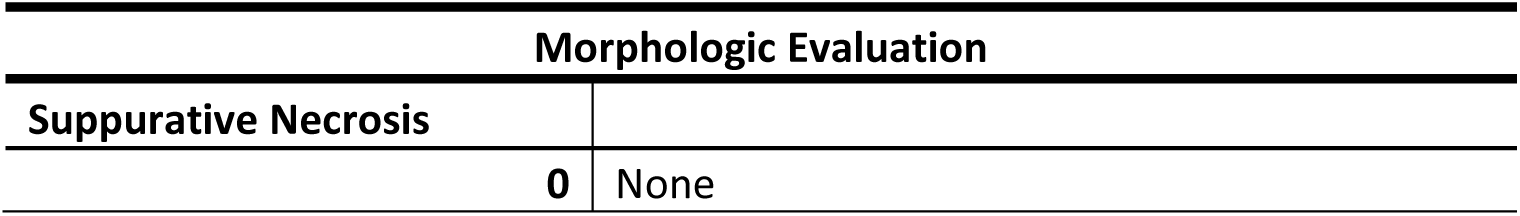
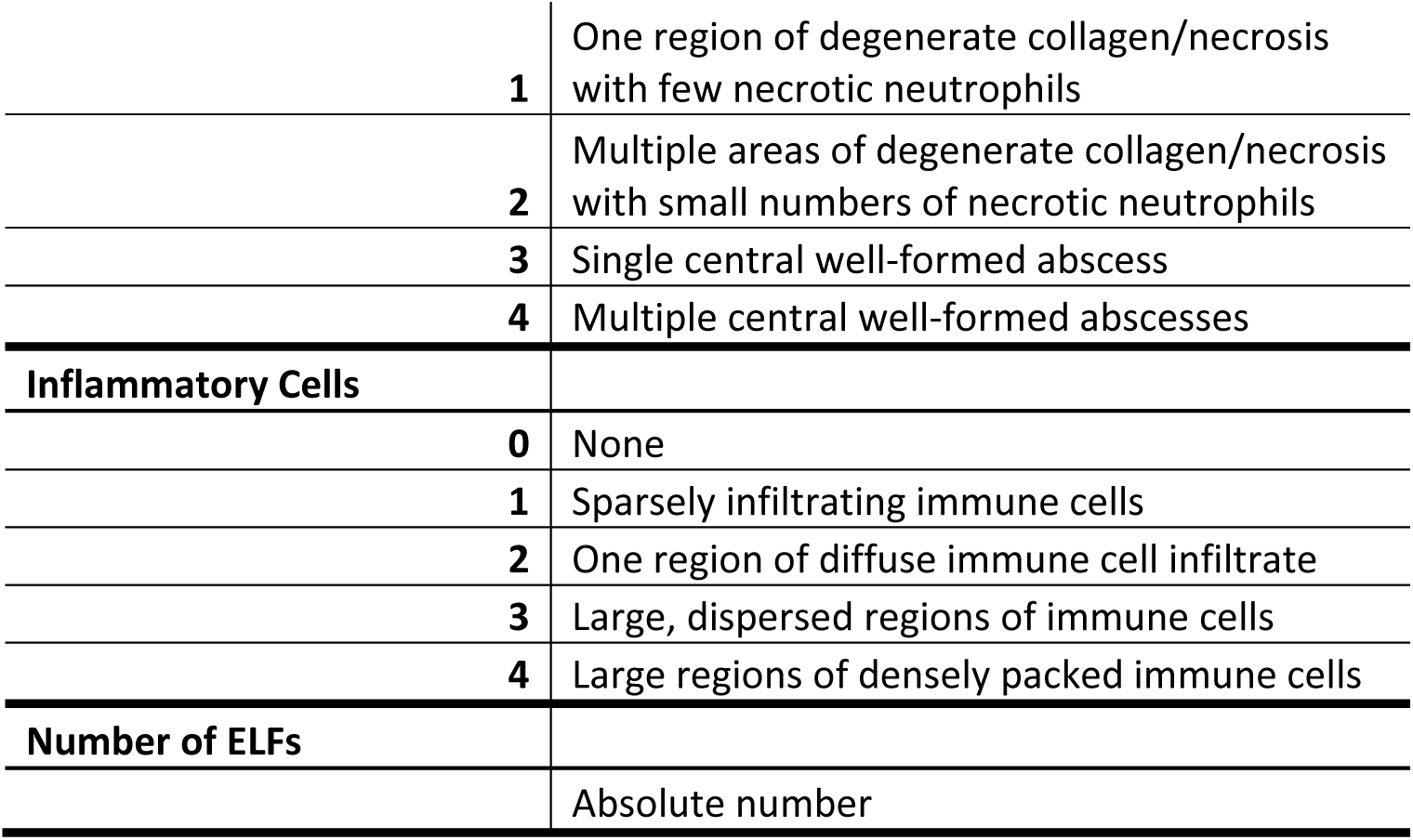
Semi-quantitative and quantitative evaluation for evaluation of histopathology of vaccine site reactions on HE-stained slides. Evaluation is divided into regions of suppurative necrosis, degree of histiocytic inflammation, and number of ectopic lymphoid follicles (ELFs).

| Morphologic Evaluation |  |
| --- | --- |
| Suppurative Necrosis |  |
| 0 | None |

|  |  |  |
| --- | --- | --- |
|  | 1 | One region of degenerate collagen/necrosis with few necrotic neutrophils |
|  | 2 | Multiple areas of degenerate collagen/necrosis with small numbers of necrotic neutrophils |
|  | 3 | Single central well-formed abscess |
|  | 4 | Multiple central well-formed abscesses |
| **Inflammatory Cells** |  |  |
|  | 0 | None |
|  | 1 | Sparsely infiltrating immune cells |
|  | 2 | One region of diffuse immune cell infiltrate |
|  | 3 | Large, dispersed regions of immune cells |
|  | 4 | Large regions of densely packed immune cells |
| **Number of ELFs** |  |  |
|  |  | Absolute number |

### Transmission Electron Microscopy

Regions within vaccination sites which were positive for anti-*C. burnetii* antibodies were extracted from formalin-fixed paraffin blocks for transmission electron microscopy as previously described (47). Briefly, tissues were cut from paraffin blocks, de-paraffinized and re-hydrated in xylene and decreasing concentrations of ethanol, then fixed in 2% glutaraldehyde and 1% osmium tetroxide. Tissues were then stained with saturated uranyl acetate and dehydrated in increasing concentrations of ethanol, followed by embedding with Eponate-12 Resin (Ted Pella, Redding, CA, USA) and cut into 100 nm sections.

### Flow Cytometry

Vaccine sites were collected and processed into single cell suspensions as previously described (29) Briefly, the skin, including the underlying skeletal muscle, at the site of elicitation was collected using a 10 mm punch biopsy and the epidermis was removed using a razor blade. The remaining tissue was placed in a gentleMACS C-tube and processed using the gentleMACS Mouse Adipose Tissue Dissociation Kit and the gentleMACS Octo Dissociator with heaters (Miltenyi Biotec). Cell suspensions were diluted to 10^7^ cells/mL and kept on ice. Cells were stained with anti-mouse CD16/CD32 for 10 min, followed by 2 µL/mL Zombie Aqua live/dead dye (Biolegend) for 5 min, and lastly with fluorochrome-conjugated cell surface antibodies for 30 min. Cells were then fixed in 4% paraformaldehyde for 20 min and kept at 4°C until analysis. Flow cytometric antibodies include CD45-APC, CD3ε-APC/Cy7, CD19-PE/Cy7, CD11b-PE, CD11c-BV711, Ly6G-FITC (Biolegend) and CD4-PerCP/Cy5.5 and CD8a-PAC Blue (BD Biosciences). Cells were evaluated using a BD LSRFortessa X-20 Flow Cytometer and analyzed using FlowJo v10.7.2 (FlowJo LLC.).

### Statistics

Statistical analyses were calculated using Prism v7.0 (GraphPad Software Inc.). Results were compared using one-way ANOVA with Dunnett’s correction for multiple comparisons as previously described (16, 24, 37). Differences were considered significant if p-value ≤ 0.05 (*), ≤ 0.01 (**), ≤ 0.001 (***), or ≤ 0.0001 (****).

### Figures

All figures were prepared using Biorender.

## Acknowledgements

We would like to thank the Dr. Malea Murphy and Texas A&M University Health Science Center Integrated Microscopy and Imaging Laboratory (IMIL) for assistance in generating the histology images and Robbie Moore and the Texas A&M School of Medicine Analytical Cytometry Core (SMACC) for assistance with the flow cytometric analysis. This work was supported by HDTRA1-14-C-0113 (JES), R01AI090142 (JES), Institutional Training Grant T32 5 OD11083-11 (APF) and Wofford Cain Endowed Research Fund (JES).

